# Vav1 is essential for HIF-1α activation in vascular response to ischemic stress

**DOI:** 10.1101/727677

**Authors:** Yongfen Min, Jaewoo Hong, Todd Wuest, P. Charles Lin

## Abstract

Vascular response to hypoxia is a major determinant of organ function under stress, which is particularly critical for vital organs such as the heart. This study identifies Vav1 as a key vascular regulator of hypoxia. Vav1 is present in vascular endothelium and is essential for HIF-1 activation under hypoxia. It regulates HIF-1α stabilization through the p38/Siah2/PHD3 pathway. Consequently, Vav1 deficient mice are predisposed to sudden death under cardiac ischemia with increased coronary endothelial apoptosis. Moreover, Vav1 binds to VEGFR1 that carries Vav1 to lysosomes for degradation in normoxia. Hypoxia upregulates Vav1 through inhibition of protein degradation. These findings reveal that regulation of Vav1 by hypoxia is analogous to HIF-1α regulation. Both proteins are constitutively produced allowing for rapid responses when stress occurs, and constantly degraded in normoxia. Hypoxia stabilizes Vav1, which is required for HIF-1α accumulation. Together they mediate the vascular response to hypoxia and maintain tissue homeostasis.

## Introduction

The vascular response to hypoxia is an important mechanism that maintains organ functions under stress, which is particularly important for vital organs such as the heart. Myocardial perfusion is a key component of cardiac homeostasis, and failure to induce sufficient perfusion represents a major cause of myocardial dysfunction and heart failure. The cellular response to hypoxia is regulated by the hypoxia-inducible factor-1 (HIF-1) transcription factor. Hypoxia stabilizes HIF-1α, allowing it to activate transcription and mediate the adaptive response. Endothelial expression of HIF-1 regulates endogenous VEGF expression and autocrine VEGF signaling is essential for endothelial cell survival ^2,3^. VEGF stimulates cellular responses by binding to cell surface receptors, VEGFR1 and VEGFR2, on vascular endothelium. VEGFR2 appears to mediate almost all of the known cellular responses to VEGF ^4^, and VEGFR1 is considered as an inhibitory receptor that acts as a decoy receptor, competing with VEGFR2 for binding to VEGF ^4^.

The levels of proteins are determined not only by synthesis, but also by degradation. Many rapidly degraded proteins function as regulatory molecules. The rapid turnover of these proteins is necessary to allow their levels to change quickly in response to external stimuli. In eukaryotic cells, two major pathways mediate protein degradation, the ubiquitin-proteasome pathway and lysosomal proteolysis, ^5^. Lysosomes are membrane-enclosed organelles that contain digestive enzymes including proteases, and these enzymes are activated by the highly acidic pH of the lysosome ^7^.

Vav1 is a guanine nucleotide exchange factor (GEF) that activates small Rho GTPase. The Vav family has three members in vertebrates with Vav1 mostly restricted to hematopoietic cells ^9,10^. Vav1 is also an important signal transducer with a pivotal role in hematopoietic cell activation, cell growth and differentiation^9-11^. Mice without Vav1 are viable, fertile and grossly normal, except for a partial block in lymphocyte development^12-14^.

This study reveals a non-hematopoietic function of Vav1, which is constantly produced and constantly undergoes protein degradation in endothelial cells in normoxia. Hypoxia blocks Vav1 degradation, and Vav1 is essential for HIF-1α stabilization. Accordingly, Vav1 deficient mice were unable to respond to hypoxic stress and exhibited a susceptibility to sudden death when subjected to acute cardiac ischemia, due to a dramatic increase of endothelial apoptosis.

## Results

### Vav1 is specifically expressed in coronary endothelium and protects mice from the development of sudden death under cardiac ischemia

Based on our previous observation of Vav1 expression in vascular endothelium {DeBusk, 2010 #25}^15^, we studied its function in vascular biology. We first validated this observation by analyzing Vav1 expression in FACS-sorted (CD31+CD45-) pulmonary microvascular endothelial cells from age and sex matched WT and Vav1 null mice (Figure 1A). We confirmed this finding by immunofluorescent staining for CD31 and Vav1 in WT mouse heart tissues. It is clear that Vav1 is specifically expressed in coronary endothelium, but not in myocardiocyte (Figure 1B).

**Figure 1.**
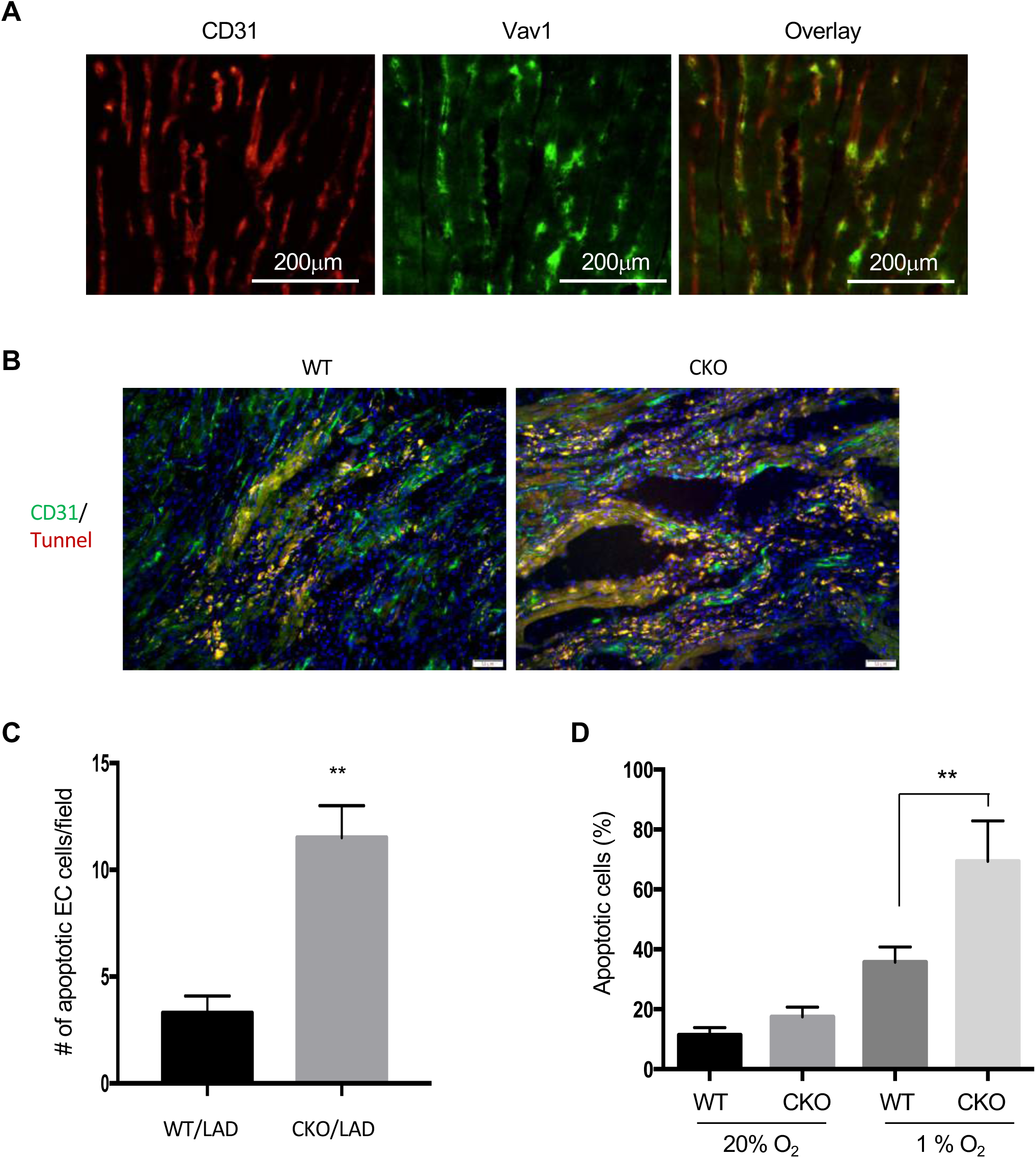
Vav1 is specifically expressed in coronary endothelium and positively regulates endothelial survival under hypoxic stress. Heart tissue sections from WT mice were subjected to immunofluorescent double staining with antibodies against CD31 (red) and Vav1 (green) (Panel A). Twelve week old sex matched WT and Vav1 endothelial specific conditional knockout mice (CKD) were subjected to LAD ligation. Heart tissues from WT and Vav1 CKD mice were harvested 2 days after LDA. Tissue sections around the ischemic region were co-stained with antibodies against either CD31 (green) or Tunnel (red) (Panel B). Representative images were shown. Apoptotic endothelial cells (double positive) were counted in ten randomly selected microscopy fields (Panel C). n= 6 mice per group, **p< 0.01. Pulmonary microvascular endothelial cells were isoalted from WT and Vav1 CKO mice, and cultured in serum free media in either normoxia (20% O2) or hypoxia (1% O2) for 12 hours, followed by flow cytometry analysis for Annexin V and 7AAD double positive apoptotic cells (Panel D). **p<0.01. The experiment was done in triplicate and repeated twice.

Myocardial ischemia occurs when blood flow to heart muscle is decreased by a blockage of coronary arteries, which is a major cause of myocardial dysfunction and heart failure. To explore the significance of Vav1 in cardiac ischemia, we employed a myocardial infarction model by ligating the left anterior descending coronary artery (LAD) to create acute ischemia in sex matched 12 week old WT and Vav1 null mice, at which time there was no significant difference in heart size and left ventricular contractility between the two groups of mice, which is in agreement with a published report^20^. Strikingly, while all the WT mice survived, all the Vav1 null mice died within hours or a day after the LAD (Figure 1C).

Since Vav1 is restricted to coronary endothelium and endothelial cell survival is a critical determinant of vascular function, we examined cell survival. A few hours after the ligation, the ischemic region of the heart tissues was harvested and subjected to double staining with CD31 and an apoptotic marker. Notably, there were significantly more double-positive apoptotic endothelial cells in heart samples collected from Vav1 null mice than WT controls (Figure 1D and 1E). To further establish the role of Vav1 in endothelial cell survival under ischemia, we isolated pulmonary microvascular endothelial cells from WT and Vav1 null mice and cultured them in either normoxic or hypoxic conditions. Consistent with the *in vivo* mouse data, deletion of Vav1 significantly increased apoptosis especially in hypoxic conditions compared to WT cells (Figure 1F). These findings reveal a critical function of Vav1 in vascular survival under hypoxia. Deletion of Vav1 in mice generates a fragile vascular phenotype, which leads to increased endothelial apoptosis and disruption of cardiac perfusion, that likely contributes to heart failure and sudden death.

### Vav1 is essential for HIF-1α stabilization *via* regulation of p38 activation in response to hypoxia

HIF-1 is a central regulator of the response to hypoxic conditions. Hypoxia stabilizes HIF-1α and activates transcription of numerous target genes. Therefore, we examined HIF-1α expression in ischemic heart samples by immuno-fluorescent staining. As expected, cardiac ischemia after LDA increased vascular HIF-1α accumulation in tissues from WT mice. Remarkably, deletion of Vav1 dramatically reduced the accumulation of HIF-1α in the vasculature (Figure 2A). To corroborate the importance of Vav1 in hypoxia induced HIF-1α accumulation, we challenge the mice with an injection of CoCl_2_ to mimic hypoxia for 5 hours. CoCl_2_ induced a strong accumulation of HIF-1α in WT heart tissues, but it completely failed to induce HIF-1α in mouse heart deficient of Vav1 (Figure 2B).

**Figure 2.**
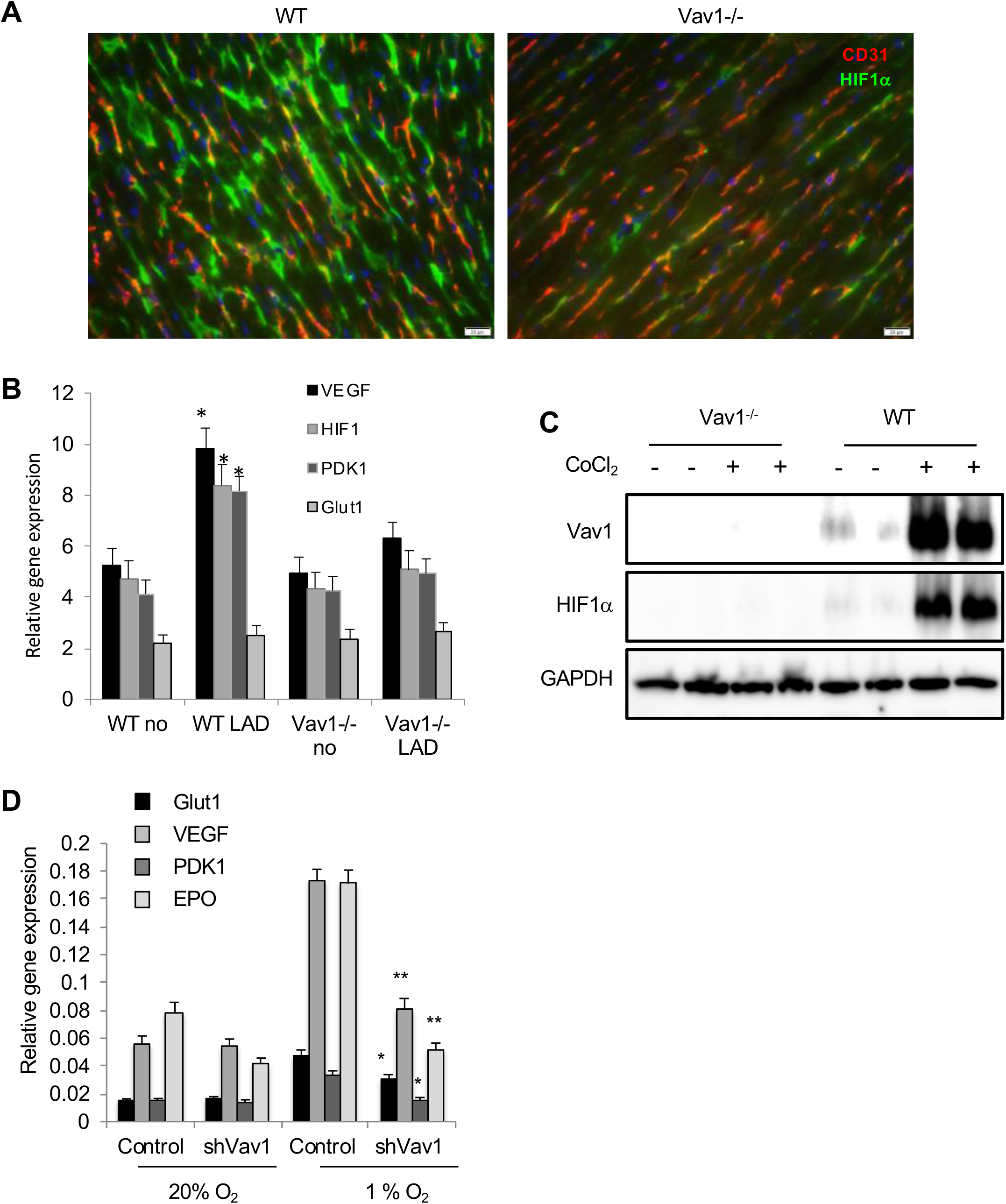
Vav1 is essential for HIF-1α induction under hypoxia. Heart tissues from the ischemic region of WT and Vav1 CKO mice were harvested 5 hours after LDA. Tissue sections were co-stained with antibodies against either CD31 (red) or HIF-1 (green) (Panel A). Representative images were shown. The ischemic region of the heart tissues were harvested from WT and Vav1 CKO mice 5 hours after LAD and subjected to qPCR for HIF-1 target gene expression (Panel B). Age and sex matched WT and Vav1 null mice received an IP injection of CoCl2 at 60mg/kg or vehicle control. Heart tissues were harvested 5 hours later and subjected to Western blot analysis for Vav1 and HIF-1a (Panel C). HUVECs were transfected with control vector or shVav1 vector for 24 hours, followed by incubation either in normoxia or hypoxia for another 24 hours. HIF-1 target gene expression was analyzed by qRT-PCR (Panel D). *p<0.05 and **p<0.01 compared to corresponding control transfected cells in hypoxia. Each experiment was done in triplicate and repeated 3 times.

To further explore the link between Vav1 and HIF-1, we knocked down Vav1 in cultured HUVECs, followed by incubation of the cells in normoxic or hypoxic conditions. The expression of HIF-1 target genes, including VEGF, EPO, Glut1 and PDK1, were analyzed by qPCR. The knockdown of Vav1 in human endothelial cells impaired the expression of HIF-1 target genes, which is particularly evident under hypoxia (Figure 2C). Collectively, these data reveal an essential role of Vav1 in hypoxia-induced HIF-1α stabilization, which drives VEGF expression in endothelial cells. Because endogenous VEGF and autocrine VEGF signaling is essential for endothelial survival^3^, these findings explain the sensitive and fragile nature of vasculature in Vav1 null mice under ischemia.

To dissect the signaling mechanism by which Vav1 regulates HIF-1α accumulation, we investigated the role of Vav1 in p38 activation, as hypoxia activates p38 ^26^ and p38 stabilizes HIF-1α ^27^. Consistent with published data, hypoxia induced p38 phosphorylation and accumulation of HIF-1α protein in HUVECs. These responses were largely blocked after a knockdown of Vav1 in these cells (Figure 3A). Conversely, ectopic expression of Vav1 further increased hypoxia-induced p38 phosphorylation as well as HIF-1α protein accumulation in hypoxia (Figure 3B). To further establish the role of p38 in Vav1 mediated HIF-1α expression, we ectopically expressed Vav1 in HUVECs and incubated the cells in hypoxia in the presence or absence of a p38 specific inhibitor, SB203580. Blocking p38 activation blunted Vav1 mediated HIF-1α accumulation (Figure 3C), which implies that p38 is downstream of Vav1, and p38 activation is required for the induction of HIF-1α accumulation upon hypoxic stimulation.

**Figure 3.**
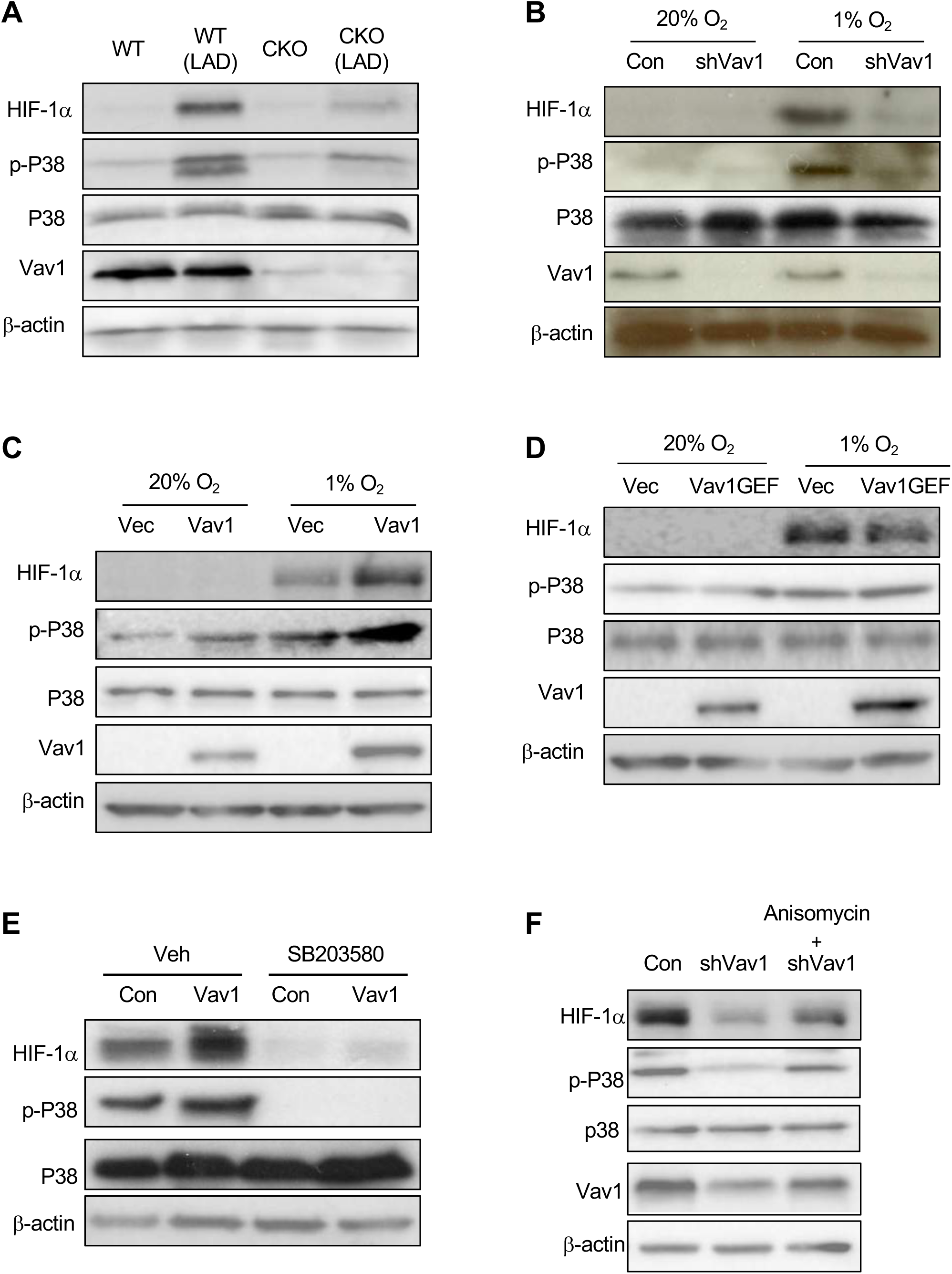
Vav1 regulates HIF-1α accumulation through p38. The heart tissues were collected from WT and Vav1 CKO mice with and without LAD, and subjected to Western blot for p35 phosphorylation Control vector or shVav1 vector transfected HUVECs were incubated under normoxia or hypoxia conditions for 24 hours. The levels of phospho p38, HIF-1a, p38 and Vav1 were analyzed by Western blot (Panel A). Identical procedures and measurements as in Panel A were done except the cells were transfecting with a control vector or Vav1 expression vector (Panel B). Control or Vav1 expression vector transfected HUVECs were incubated in the absence or presence of SB203580 at 10 μM in hypoxia for 24 hours. The levels of HIF-1a, p38 and phospho-p38 were analyzed by Western blot (Panel C). Control vector or shVav1 vector transfected HUVECs were incubated under normoxic or hypoxic conditions for 24 hours. The levels of phospho HIF-1a, Vav1, P-p38, p38, P-siah2, Siah2 and PHD3 were analyzed by Western blot (Panel D). Each experiment was repeated 3 times.

Next, we examined phosphorylation of Siah2 as p38 directly phosphorylates this protein that targets prolyl hydroxylase-3 (PHD3) for degradation ^22,23^. We found that hypoxia activated p38, which correlated with increased phosphorylation of Siah2 and reduction of PHD3 in vector transfected cells (Figure 3D). In contrast, knockdown of Vav1 inhibited hypoxia-mediated phosphorylation of p38 and Siah2, which correlated with an increase of PHD3 (Figure 3D). As PHD3 targets HIF-1α for degradation, a reduction of PHD3 correlated with an increase of HIF-1α in Vav1 knockdown cells (Figure 3D). These results suggest that Vav1 regulates HIF-1α accumulation through the p38 mediated Siah2/PHD3 pathway.

### Hypoxia upregulates Vav1 levels *via* inhibition of lysosomal mediated protein degradation

Based on the role of Vav1 in regulating vascular response to hypoxia, we investigated the mechanism by which hypoxia regulates Vav1. We found that mimic hypoxia *in vivo* using CoCl_2_ induced a dramatic increase of Vav1 levels in WT heart tissues (Figure 2B). To validate the finding, we incubated HUEVCs in normoxia or hypoxia for 5 hours. Hypoxia increased Vav1 protein levels (Figure 4A), however hypoxia had no significant effect on mRNA levels of *VAV1* (Figure 4B). In addition, we knocked down HIF-1α and incubated the cells in hypoxia. Neutralization of HIF-1 had no impact on Vav1 protein levels (Figure 4C). Moreover, Vav1 level was not affected by HIF-1α downregulation by PX-478 but when Vav1 was controlled by shRNA or overexpression, the level of HIF-1α was controlled by Vav1 parallelly (Supplementary Figure 1). This set of results implies that hypoxia-mediated Vav1 expression does not occur at the transcriptional level.

**Figure 4.**
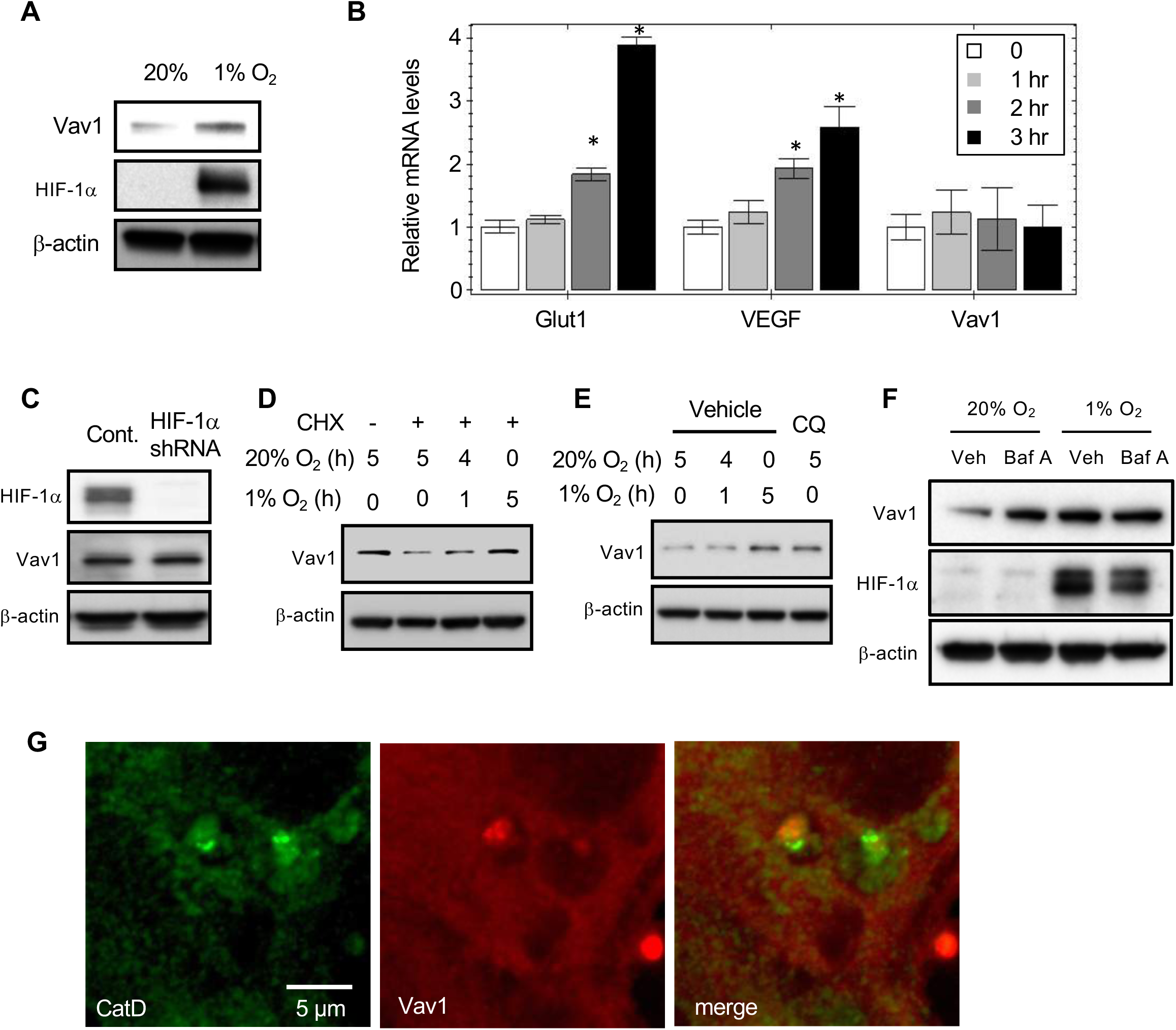
Hypoxia upregulates Vav1 through inhibition of lysosomal mediated protein degradation. HUVECs were cultured either in 20% or 1% O2 for 4 hours, followed by Western blot analysis (Panel A). qPCR analysis of Glut1, VEGF and Vav1 in HUVECs cultured in 1% O2 for 0, 1, 2, or 3 hours. *p<0.05 compared to corresponding time 0 in each group (Panel B). The levels of Vav1 and HIF-1a were measured by Western blot of HUVECs transduced with HIF-1a or scramble shRNA expressing Lentivirus for 24hrs (Panel C). HUVECs were incubated in a combination of normoxic to hypoxic environment at indicated times for a total of 5hrs in the presence or absence of cycloheximide at 10mM. The protein levels were measured by Western blot from the total lysate (Panel D). The levels of Vav1 were measured by Western blot for HUVECs either cultured in 20% O2 in the presence of chloroquine (CQ) at 50mM for 5 hours or cultured in 1% O2 for different times (Panel E). The levels of Vav1 were measured by Western blot for HUVECs cultured in either 20% or 1% O2 in the absence or presence of Bafilomycin A (Baf A) at 100nM for 5 hrs (Panel F). Immunofluorescent staining for Vav1 (red) and Cathepsin D (green) in HUVECs were imaged by a LSM780 confocal microscope (Panel G). Each experiment was repeated at least twice and representative images are shown.

Next, we investigated whether the regulation occurs via protein degradation. To test this hypothesis, HUVECs were cultured in the presence of cycloheximide to suppress nascent protein synthesis with a sequential incubation of the cells in normoxia and hypoxia for 5 hours. Interestingly, addition of cycloheximide led to a significance reduction of Vav1 protein compared to vehicle treated cells in normoxia. Incubation of the cells in hypoxia resulted in a time dependent increase of Vav1 protein in the presence of cycloheximide (Figure 4D). These findings imply that Vav1 protein is constantly produced and constantly undergoes degradation.

To distinguish if Vav1 degradation is mediated through the proteasomal or lysosomal pathway, we cultured HUVECs in normoxia in the presence of lactacystin to inhibit proteasomal pathway or chloroquine, a lysosomal inhibitor. Notably, addition of chloroquine resulted in a significant increase of Vav1. The levels of Vav1 in the presence of chloroquine under normoxia were close to the levels observed when cells were cultured in hypoxia for several hours (Figure 4E). In contrast, addition of lactacystin did not significantly affect the levels of Vav1 (data not shown).

To corroborate the role of lysosomes in Vav1 degradation, we utilized an inhibitor for lysosomal acidification, bafilomysin-A (BafA). The result confirmed that blocking lysosomal activation led to increased Vav1 in normoxia. Hypoxia upregulated Vav1, and addition of Baf A had no additional effect in hypoxia (Figure 4F). We also examined if Vav1 is present in lysosomes by performing immunofluorescent staining with antibodies against Vav1 and cathepsin D (CatD), a lysosomal marker, in HUVECs. Vav1 protein co-localized in lysosomes with CatD, and it appeared that the levels of Vav1 were lower where CatD was strong and vise verse (Figure 4G). Collectively, these results suggest that lysosomes mediate Vav1 degradation in normoxia, and hypoxia blocks protein degradation, leading to Vav1 accumulation.

### VEGFR1-bound Vav1 is carried to lysosomes

To investigate the mechanism by which Vav1 moves to lysosomes, we first examined Vav1 ubiquitination status in HUVECs as protein ubiquitination is a sorting signal that targets protein to multivesicular bodies (MVBs). Hypoxia increased total protein ubiquitination in both vector and Vav1 construct transfected cells (Suppl. Figure 2 A). However, there was no detectable ubiquitination of Vav1 in both normoxic and hypoxic groups (Suppl. Figure 2 B and C), which led us to speculate there might be a carrier that moves Vav1 to lysosomes. The four amino acids YXEP were identified as a specific binding motif recognized by the SH2 domain of Vav1 (Songyang, 1994 #81). Sequence alignment revealed the presence of this motif in VEGFR1 (aa990-993), but not in VEGFR2.

To test if Vav1 binds VEGFR1, we ectopically expressed Vav1 and VEGFR1 in HeLa cells, followed by immunofluorescent staining. Confocal microscopy analysis detected a strong FRET signal between Vav1 and VEGFR1 in MVB (Figure 5A). Subsequently, we performed a pull-down experiment in HeLa cells co-transfected with expression vectors for Vav1 (GFP tagged) and VEGFR1. Vav1 was specifically pulled-down together with VEGFR1 (Figure 5B). Lastly, we generated a deletion mutant of VEGFR1 by deleting the four amino acid Vav1-binding sequence (ΔYKEP). We co-expressed Vav1 (HA tagged) with VEGFR1 or the mutant construct, followed by pull-down analysis. The data show that only WT VEGFR1 was able to pull-down Vav1, but not ΔYKEP (Figure 5C). These results confirm direct binding of Vav1 to VEGFR1.

**Figure 5.**
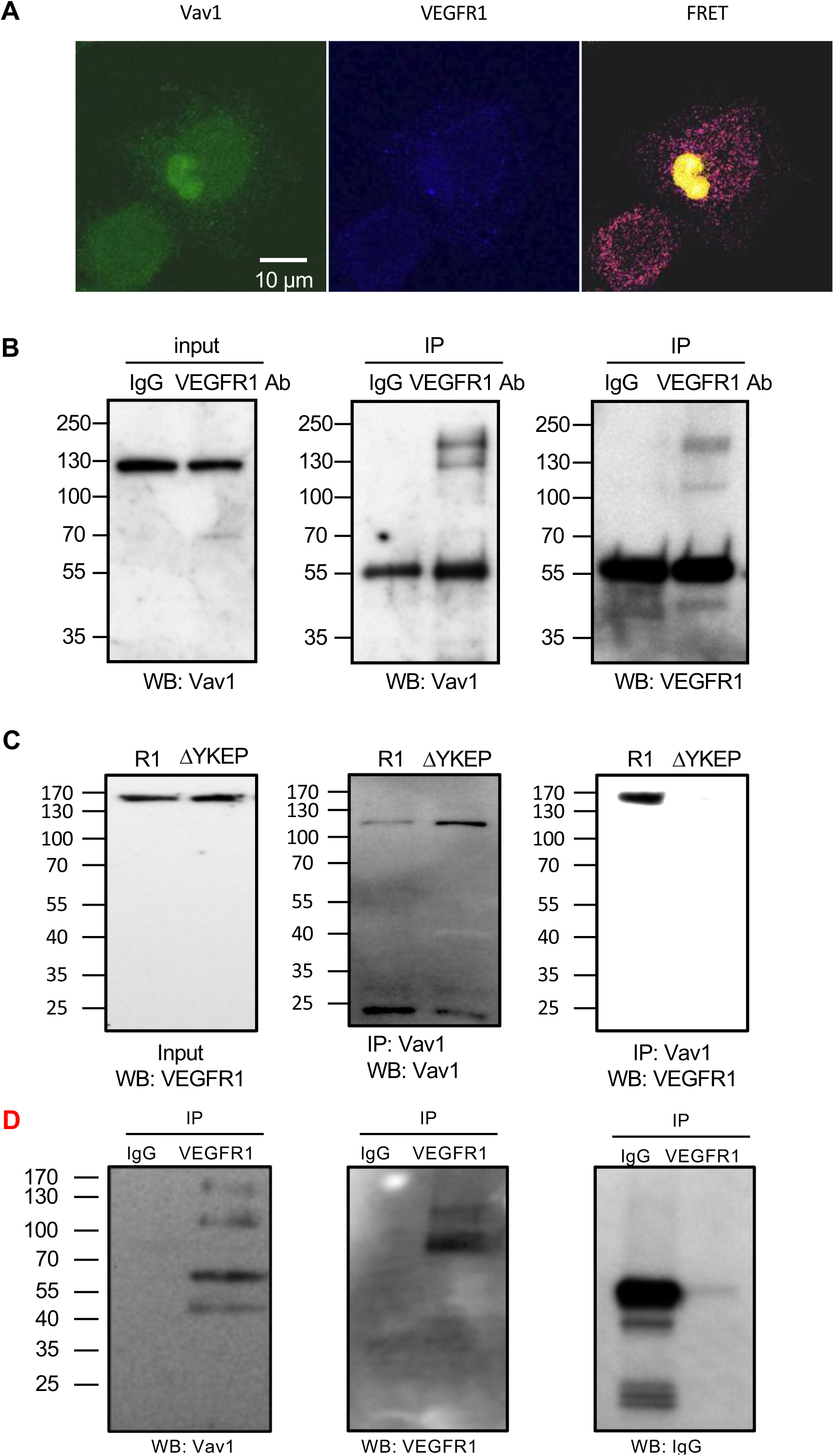
Vav1 binds to VEGFR1. HeLa cells were co transfected with expression vectors for Flag-Vav1 and VEGFR1-V5 for 24hrs, followed by FRET analysis using LSM780 confocal microscope and ImageJ (Panel A). Cell lysates from Vav1 and VEGFR1 vector transfected HeLa cells were subjected to immunoprecipitation with either VEGFR1 or control antibody. The input and pull-downs were subjected to Western blot for Vav1 or VEGFR1 (Panel B). VEGFR1 (R1) or YKEP deletion VEGFR1 (ΔYKEP) was co-expressed with Vav1 in HeLa cells. The input and pull-downs with Vav1 antibodies were analyzed by Western blot (Panel C). Cell lysates from HUVECs were subjected to immunoprecipitation with either VEGFR1 or control antibody. The input and pull-downs were subjected to Western blot for Vav1 or VEGFR1 (Panel D). Each experiment was repeated twice and representative images are shown.

Since receptor tyrosine kinases have been shown to undergo lysosomal-mediated degradation after activation, we investigated if VEGFR1 carries Vav1 to lysosomes for degradation. To test this hypothesis, we co-tranfected HUVECs with expression vectors for Vav1 and VEGFR1, followed by immunofluorescent staining for these two proteins and Lamp1, a lysosomal marker. The results show co-localization of Vav1 and VEGFR1 in lysosomes (Figure 6A). We then tested whether binding with VEGFR1 is required for Vav1 translocation to lysosomes. HUVECs were cotransfected with expression vectors for Vav1 with VEGFR1 or ΔYKEP mutant, followed by staining for Vav1, VEGFR1 and Lysobright, a marker for lysosomes. As expected, disruption of VEGFR1 binding prevented Vav1 being localization in lysosomes (Figure 6B). The data confirm that Vav1 binds to VEGFR1 and VEGFR1 carries Vav1 to lysosomes.

**Figure 6.**
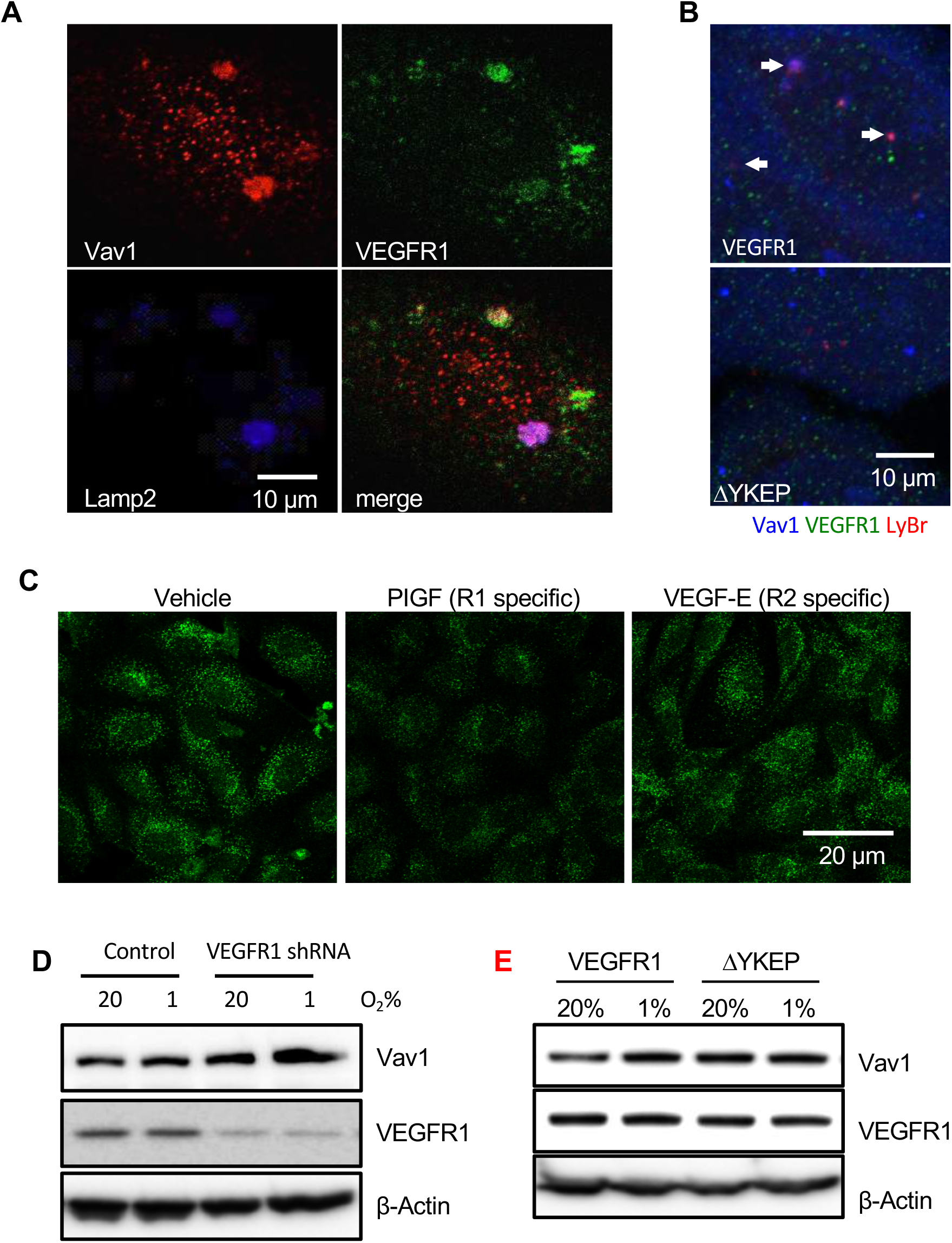
VEGFR1 carries Vav1 to lysosomes for degradation. HUVECs were immunostained for Lamp2, Vav1 and VEGFR1, and imaged under confocal microscopy (Panel A). HeLa cells were transfected with expression vectors for Vav1 with either VEGFR1 or ΔYKEP. Cells were stained with LysoBrite (red), VEGFR1 (green) and Vav1 (blue). White arrows point to colocalization of Vav1, VEGFR1 and LysoBrite (Panel B). HUVECs were stimulated with either PLGF or VEGF-E at 50ng/ml for 5 hours, followed by staining with antibodies for Vav1 and imaged under confocal microscopy (Panel C). HUVECs were infected with Lentiviral vectors for scramble or VEGFR1 shRNA and cultured in either 20% O2 or 1% O2 incubators. Cell lysate was analyzed by Western blot for Vav1 and VEGFR1 (Panel D). HUVECs were transfected with expression vectors for Vav1 with either VEGFR1 or ΔYKEP VEGFR1. The levels of Vav1 and VEGFR1 were assessed by Western blot (Panel E). Each experiment was repeated at least twice and representative images are shown.

The above findings led us to examine if activation of VEGFR1 induces Vav1 degradation. To do so, we stimulated HUVECs with PLGF, a VEGFR1 specific ligand; or VEGF-E, a VEGFR2 specific ligand, for 30 minutes. As predicted, activation of VEGFR1 induced Vav1 degradation, while activation of VEGFR2 that does not binds to Vav1 did not change the levels of Vav1 compared to controls (Figure 6C). Moreover, knockdown of VEGFR1 using shRNA in HUVECs significantly increased the levels of Vav1 in normoxia and this increase was more pronounced in hypoxia (Figure 6D). Conversely, ectopic expression of ΔYKEP led to an increase of Vav1 compared to VEGFR1 transfected cells in normoxia (Figure 6E). Hypoxia increased Vav1 levels in VEGFR1 transfected cells, but the levels of Vav1 in ΔYKEP transfected cells remained the same in normoxia and hypoxia (Figure 6E). Collectively, these data suggest that VEGFR1 carries Vav1 to lysosomes and activation of the receptor induces Vav1 degradation.

## Discussion

The vascular response to hypoxia is a powerful mechanism to maintain organ function and to reduce the negative effects that hypoxia otherwise would produce. Thus, identification of molecular mediators that regulate vascular homeostasis is of great importance. This study identifies Vav1 as a key regulator of the vascular response to hypoxia. Vav1 is continually produced and degraded in normoxic conditions. Hypoxia blocks the degradation, leading to Vav1 accumulation. Vav1 is required for hypoxia-induced HIF-1α accumulation. Consequently, deletion of Vav1 in mice prevents HIF-1 activation upon cardiac ischemic stress, which leads to endothelial apoptosis, heart failure and death of the animal.

Vav1 is considered as a hematopoietic specific protein ^9-11^. The Vav1 promoter-driven Cre mice are commonly used for specific gene deletion in hematopoietic cells. However, the current study reveals expression of Vav1 in endothelial cells. This finding is in agreement with a genetic tracing study indicating Vav1 in endothelium ^15^, as well as the notion that endothelial cells and blood cells are derived from a common progenitor, and they often share common mediators and pathways. This finding raises a concern regarding specificity when using Vav1-Cre mice for gene deletion in hematopoietic cells.

Endothelial-specific deletion of HIF-1 disrupts a hypoxia driven VEGF autocrine loop ^2^. Endogenous production of VEGF in endothelial cells and cell-autonomous activity is crucial for vascular homeostasis. In the absence of autocrine VEGF signaling, endothelial cells undergo apoptosis ^3^. This phenotype is manifested without detectable changes in the total levels of VEGF and cannot be rescued by exogenous VEGF ^3^. Our data demonstrate that Vav1 is essential for HIF-1α accumulation in hypoxia. In the absence of Vav1, the endothelium is unable to achieve HIF-1 activation and induction of VEGF, leading to a significant increase in endothelial apoptosis under stress.

Hypoxia activates p38 MAPK ^26^, and p38 stabilizes HIF-1α ^27^. In T cells, Vav1 acts as a point of integration of signal transduction for receptor-mediated p38 activation^21^. Consistent with these findings, we show that hypoxia induces p38 phosphorylation and HIF-1α accumulation in endothelial cells, which is totally dependent on Vav1. Without Vav1, hypoxia fails to activate p38 thereby interrupting the pathway of HIF-1α accumulation. The ubiquitin ligase Siah2 regulates the stability of prolyl hydroxylase-3 (PHD3) that targets HIF-1α for degradation ^23^. p38 phosphorylates Siah2, which increases Siah2-mediated degradation of PHD3 thus preventing HIF-1α degradation ^22^. Our data suggest that Vav1 is upstream of p38 and is essential for hypoxia-mediated activation of p38. Without Vav1, hypoxia is unable to activate p38, preventing the subsequent activation of Siah2 and PHD3 degradation, necessary for HIF-1α accumulation.

Remarkably, Vav1 is continually produced in endothelial cells and has a high turn-over rate due to lysosomal mediated degradation. These findings reveal that the regulation of Vav1 is analogous to HIF-1α regulation. Both proteins are constitutively produced and both are continually degraded via lysosomal (Vav1) and proteosomal (HIF-1α) pathways under normoxia. Hypoxia stabilizes Vav1 and Vav1 is essential for HIF-1α accumulation. Together, these two proteins are key mediators of the vascular response to hypoxia.

It has been reported that Vav1 is targeted to lysosomes for degradation in pancreatic tumor cells ^28^. Contrary to tumor cells in which Vav1 is targeted to lysosomes through interaction with the cytoplasmic chaperone Hsc70^28^, we found that Vav1 is transported to lysosomes by a carrier protein, VEGFR1, in endothelial cells. Vav1 binds to VEGFR1. Knockdown of VEGFR1 inhibits Vav1 degradation, and conversely activation of VEGFR1 increases Vav1 degradation. This finding provides a new mechanism by which VEGFR1 inhibits VEGF mediated angiogenesis. Activation of this receptor induces Vav1 degradation, a GEF protein for small RhoGTPase, and thus plays a negative role in cell motility and angiogenesis.

In summary, this study reports a protective role of Vav1 in vascular biology. Hypoxia upregulates Vav1 and Vav1 is essential for HIF-1α accumulation and vascular response to stress. Vav1 null mice exhibit a markedly reduced capacity to survive the stress of acute myocardial infarction. As sudden death in the setting of myocardium infarct is a very important medical issue, our findings could have important implications in the etiology and treatment of this pathologic condition.

## Materials and methods

### Animals

The mice were maintained in a pathogen-free facility at the National Cancer Institute (Frederick, MD) in accordance with Animal Care and Use Committee regulations. C57BL/6J mice were purchased from the Jackson Lab and Vav1 null mice on the C57/BL6 background were kindly provided by Dr. Victor Tybulewicz at the MRC National Institute for Medical Research, UK^14^. Sex and age-matched mice were used in all the studies. The LAD ligation was performed as described ^29^. Heart function was evaluated by Echocardiography.

### Cell Culture and Reagents

HUVECs (Lonza, Walkersville, MD) were cultured with EGM-2 medium (Lonza), and maintained at 37°C with 5% CO_2_. Hypoxia was performed by incubating cells in an incubator with 1% O_2_ (Thermo, Middletown, VA). Human PLGF and VEGF-E were purchased from ProSpec (East Brunswick, NJ). Cycloheximide and chloroquine were purchased from Tocris (Bristol, UK), bafilomysin-A (BafA) and CoCl_2_ were from Sigma. HeLa cells were cultured in DMEM medium.

### Immunohistochemistry

Frozen tissue sections were incubated with anti-CD31, anti-Vav1 or anti HIF-1α antibody (Pharmingen). Apoptotic cells were evaluated by the ApopTag® Red In Situ Apoptosis Detection Kit (Chemicon) according to manufacturer’s instructions. The number of apoptotic endothelial cells were calculated by counting CD31 and TUNEL double positive cells in ten randomly selected high power fields under microscopy.

### Western blot and immunoprecipitation

Endothelial cell lysates from WT and Vav1 null mice were subjected to western blot and incubated with anti Vav1 antibody. For hypoxic treatment, HUVECs were incubated under either 20% O_2_ or 1% O_2_ for 24 hours. The levels of HIF-1α, Siah2, pSiah2, PHD3, p38 and phospho-p38 were analyzed by Western blot using specific antibodies (Cell Signaling, Danvers, MA). SB 203580 (Cell Signaling) at 10 μM was used to inhibit p38 phosphorylation.

For immunoprecipitation, cells were lysed with lysis buffer (Cell Signaling) and immunoprecipitated with antibodies against Flag (Sigma, St. Louis, MO) or VEGFR1 (Genetex, Irvine, CA) overnight followed by protein A/G magnetic beads (Pierce, Waltham, MA). The membranes were probed with antibodies against Vav1 (EMD Millipore, Billerica, MA), Flag, HIF1α (BD Biosciences, San Jose, CA), VEGFR1 (Abcam, Cambridge, UK) and Ubiquitin (Cell Signaling).

### Real-Time RT-PCR analysis

Real time RT-PCR was performed using total RNA isolated on RNeasy Quick spin columns (QIAGEN, CA). One μg of total RNA was used to perform reverse transcriptase–polymerase chain reaction (RT-PCR) using iScript supermix (Biorad, Hercules, CA). The sequence of PCR primers used are: Vav1, 5’-CAACCTGCGTGAGGTCAAC −3’ and 5’-ACCTTGCCAAAATCCTGCACA −3’ VEGF, 5’-TGTACCTCCACCATGCCAAGT-3’ and, 5’-CGCTGGTAGACGTCCATGAA-3’; PDK1, 5’-ACCAGGACAGCCAATACAAG-3’, and 5’-CCTCGGTCACTCATCTTCAC-3’; Glut1, 5’-ACGCTCTGATCCCTCTCAGT-3’ and 5’-GCAGTACACACCGATGATGAAG-3’; EPO, 5’-ACCAACATTGCTTGTGCCAC-3’ and 5’-TCTGAATGCTTCCTGCTCTGG-3’. Values are expressed as fold increase relative to the reference sample (untreated control) and analyzed with CFX manager (Biorad). All primers were purchased from Sigma.

### Immunofluorescent Staining and Fluorescent Resonance Energy Transfer (FRET)

HeLa was transfected with Flag-tagged Vav1 and V5-tagged VEGFR1 grown on coverslips were stained with antibody against Flag-tag (Sigma), and against V5-tag (Cell Signaling). Confocal microscopy was performed with LSM-780 (Zeiss, Oberkochen, Germany) and analyzed with ImageJ (NIH, Bethesda, MD).

### Statistical analysis

All statistical analyses were carried out using Prism 6 (La Jolla, CA). Quantitative variables were analyzed by t-test, one-way ANOVA test. All statistical analysis were two-sided, and p<0.05 was considered statistically significant.

## Acknowledgements

This research was supported by the Intramural Research Program of the Center for Cancer Research, National Cancer Institute, National Institutes of Health. We thank Dr. Stephen Anderson in the NCI for proof reading the manuscript.

## References

1. Hill, J.A. & Olson, E.N. Cardiac plasticity. N Engl J Med 358, 1370–1380 (2008).

2. Tang, N., et al. Loss of HIF-1alpha in endothelial cells disrupts a hypoxia-driven VEGF autocrine loop necessary for tumorigenesis. Cancer Cell 6, 485–495 (2004).

3. Lee, S., et al. Autocrine VEGF signaling is required for vascular homeostasis. Cell 130, 691–703 (2007).

4. Ferrara, N., Gerber, H.P. & LeCouter, J. The biology of VEGF and its receptors. Nat Med 9, 669–676 (2003).

5. Ciechanover, A. Proteolysis: from the lysosome to ubiquitin and the proteasome. Nat Rev Mol Cell Biol 6, 79–87 (2005).

6. Etlinger, J.D. & Goldberg, A.L. A soluble ATP-dependent proteolytic system responsible for the degradation of abnormal proteins in reticulocytes. Proc Natl Acad Sci U S A 74, 54–58 (1977).

7. Hicke, L. & Riezman, H. Ubiquitination of a yeast plasma membrane receptor signals its ligand-stimulated endocytosis. Cell 84, 277–287 (1996).

8. Schneider, D.L. ATP-dependent acidification of intact and disrupted lysosomes. Evidence for an ATP-driven proton pump. J Biol Chem 256, 3858–3864 (1981).

9. Tybulewicz, V.L., Ardouin, L., Prisco, A. & Reynolds, L.F. Vav1: a key signal transducer downstream of the TCR. Immunol Rev 192, 42–52 (2003).

10. Bustelo, X.R. Regulatory and signaling properties of the Vav family. Mol Cell Biol 20, 1461–1477 (2000).

11. Turner, M. & Billadeau, D.D. VAV proteins as signal integrators for multi-subunit immune-recognition receptors. Nat Rev Immunol 2, 476–486 (2002).

12. Fischer, K.D., et al. Defective T-cell receptor signalling and positive selection of Vav-deficient CD4+ CD8+ thymocytes. Nature 374, 474–477 (1995).

13. Zhang, R., Alt, F.W., Davidson, L., Orkin, S.H. & Swat, W. Defective signalling through the T- and B-cell antigen receptors in lymphoid cells lacking the vav proto-oncogene. Nature 374, 470–473 (1995).

14. Turner, M., et al. A requirement for the Rho-family GTP exchange factor Vav in positive and negative selection of thymocytes. Immunity 7, 451–460 (1997).

15. Georgiades, P., et al. VavCre transgenic mice: a tool for mutagenesis in hematopoietic and endothelial lineages. Genesis 34, 251–256 (2002).

16. Walsh, K. & Shiojima, I. Cardiac growth and angiogenesis coordinated by intertissue interactions. J Clin Invest 117, 3176–3179 (2007).

17. Shiojima, I., et al. Disruption of coordinated cardiac hypertrophy and angiogenesis contributes to the transition to heart failure. J Clin Invest 115, 2108–2118 (2005).

18. Tirziu, D., et al. Myocardial hypertrophy in the absence of external stimuli is induced by angiogenesis in mice. J Clin Invest 117, 3188–3197 (2007).

19. Heineke, J., et al. Cardiomyocyte GATA4 functions as a stress-responsive regulator of angiogenesis in the murine heart. J Clin Invest 117, 3198–3210 (2007).

20. Sauzeau, V., Jerkic, M., Lopez-Novoa, J.M. & Bustelo, X.R. Loss of Vav2 proto-oncogene causes tachycardia and cardiovascular disease in mice. Mol Biol Cell 18, 943–952 (2007).

21. Hehner, S.P., Hofmann, T.G., Dienz, O., Droge, W. & Schmitz, M.L. Tyrosine-phosphorylated Vav1 as a point of integration for T-cell receptor- and CD28-mediated activation of JNK, p38, and interleukin-2 transcription. J Biol Chem 275, 18160–18171 (2000).

22. Khurana, A., et al. Regulation of the ring finger E3 ligase Siah2 by p38 MAPK. J Biol Chem 281, 35316–35326 (2006).

23. Nakayama, K., Qi, J. & Ronai, Z. The ubiquitin ligase Siah2 and the hypoxia response. Mol Cancer Res 7, 443–451 (2009).

24. Zakaria, S., et al. Differential regulation of TCR-mediated gene transcription by Vav family members. J Exp Med 199, 429–434 (2004).

25. DeBusk, L.M., Boelte, K., Min, Y. & Lin, P.C. Heterozygous deficiency of delta-catenin impairs pathological angiogenesis. J Exp Med 207, 77–84 (2010).

26. Kulisz, A., Chen, N., Chandel, N.S., Shao, Z. & Schumacker, P.T. Mitochondrial ROS initiate phosphorylation of p38 MAP kinase during hypoxia in cardiomyocytes. Am J Physiol Lung Cell Mol Physiol 282, L1324–1329 (2002).

27. Emerling, B.M., et al. Mitochondrial reactive oxygen species activation of p38 mitogen-activated protein kinase is required for hypoxia signaling. Mol Cell Biol 25, 4853–4862 (2005).

28. Razidlo, G.L., et al. Dynamin 2 potentiates invasive migration of pancreatic tumor cells through stabilization of the Rac1 GEF Vav1. Dev Cell 24, 573–585 (2013).

29. Bialik, S., et al. Myocyte apoptosis during acute myocardial infarction in the mouse localizes to hypoxic regions but occurs independently of p53. J Clin Invest 100, 1363–1372 (1997).

