## Supplementary material for "Vav1 is essential for HIF-1α activation in vascular response to ischemic stress": suppl fig

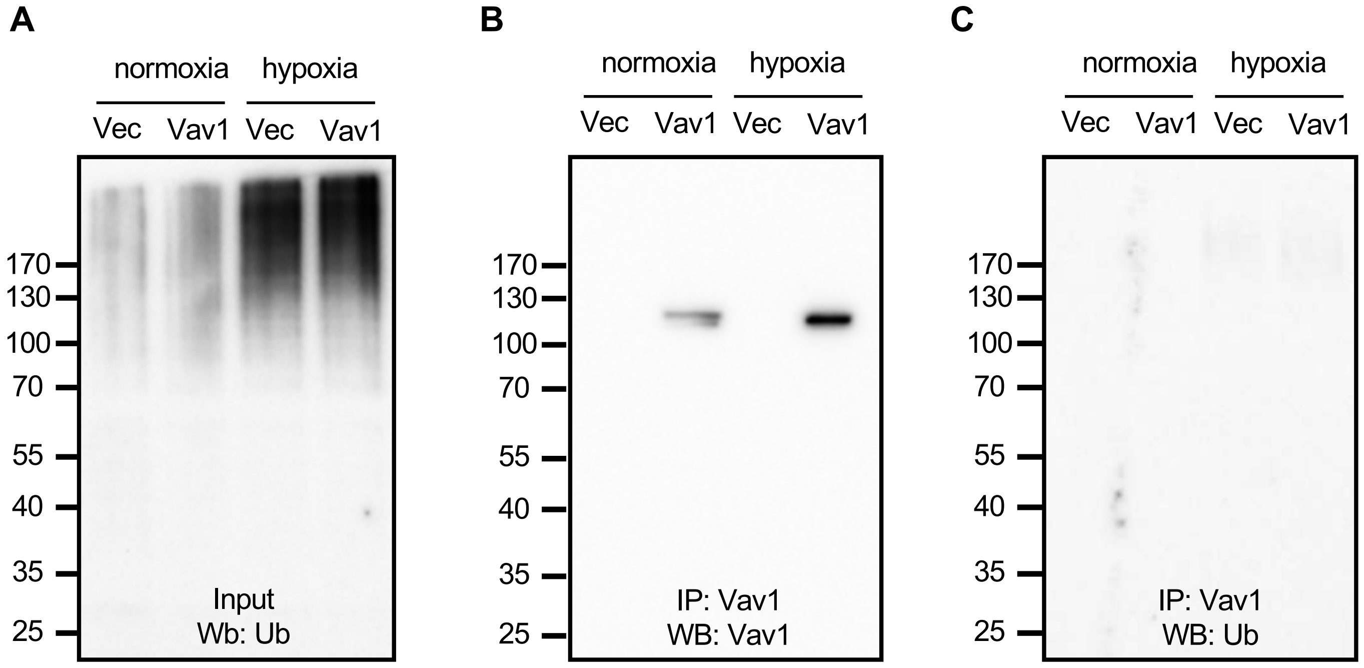


**Supplementary Figure 1**. Vav1 is not ubiquitinated in endothelial cells. HUVECs were infected with either lentiviral empty vector or Vav1 expressing vector for 48 hours, followed by an incubation in either normoxic or hypoxic conditions for an additional 5 hours. The total ubiquitin levels were measured from the input (Panel A). The Vav1 was immunoprecipitated from the cell lysate and probed for Vav1 (Panel B) and ubiquitin (Panel C) by Western blot. Each experiment was repeated twice.
