## Supplementary material for "Vav1 is essential for HIF-1α activation in vascular response to ischemic stress": suppl legend

Supplementary Figure 1. Vav1 is not ubiquitinated. Empty vector or Vav1 overexpressing lentivirus was transduced to HUVECs. 48 hours after transduction, cells were incubated in normoxic or hypoxic incubator for 5 hours and then Vav1 was pulled down from the total lysate of the transduced cells. The total ubiquitin level was measured from the input lysate (A). The Vav1 (B) and ubiquitin (C) levels were measured from the pull down products in order to detect whether ubiquitin is bound to Vav1.
